# Liquid-liquid phase separation and aggregation of a globular folded protein SUMO1

**DOI:** 10.1101/2023.06.10.544456

**Authors:** Simran Arora, Debsankar Saha Roy, Sudipta Maiti, Sri Rama Koti Ainavarapu

## Abstract

Many studies in recent years have investigated the phenomenon of liquid-liquid phase separation (LLPS) in proteins. LLPS is reported in intrinsically disordered proteins (IDPs) or in proteins with intrinsically disordered regions (IDRs) lacking a well-defined three-dimensional structure. However, the occurrence of LLPS in folded proteins, that lacks IDRs is not widely known. It is generally assumed that the compact structure and limited flexibility of folded proteins hinder their ability to establish weak and dynamic interactions crucial for LLPS. Contrary to the prevailing understanding, we present direct evidence of rapid phase separation of a globular protein, SUMO1, occurring under crowded conditions at physiological pH and room temperature. The protein molecules in the liquid droplets undergo conformational changes with time, monitored by intrinsic tryptophan fluorescence. We also demonstrate the phase transition of SUMO1 droplets from liquid to solid state with maturation, ultimately leading to aggregation. The SUMO1 aggregates contain a significant amount of beta-sheet structure but have amorphous morphology, probed by several spectroscopic techniques (Thioflavin T fluorescence, Raman Spectroscopy, and TEM). Our findings provide insights into the behaviour of SUMO1 protein in crowded environment, albeit, the underlying mechanism is not well understood and may differ from those of IDPs. Furthermore, SUMO1 protein is known to colocalize in inclusion bodies of IDPs, and therefore, LLPS of SUMO1 may have important biological implications in neurodegenerative diseases.

## Introduction

The eukaryotic cellular organization encompasses a variety of membrane-bound organelles, such as mitochondria^1^ and lysosomes^2^, as well as membrane-less organelles like the nucleolus^3^ and stress granules^4,5^. Recent studies have indicated that the process of liquid-liquid phase separation (LLPS) leading to the formation of membrane-less organelles contributes to the spatial organization within cells.^6–8^ LLPS involves the formation of liquid droplets in the cytoplasm or in the nucleoplasm facilitated by diverse interactions between proteins and nucleic acids.^9^ These droplets, also referred to as bio-molecular condensates, play a crucial role in organizing the contents of living cells and also host a number of fundamental processes.^10–12^ However, LLPS is also associated with neurodegenerative diseases including amyotrophic lateral sclerosis (ALS), frontotemporal dementia (FTD), Alzheimer’s disease (AD), and Parkinson’s disease (PD).^13^ These diseases involve the aggregation of proteins to form pathological inclusions inside the cell. In recent years, it has been shown that many intrinsically disordered proteins (IDPs) such as α-synuclein^14^, FUS^15–18^, TDP-43^18–20^, and tau^21– 23^ found in these pathological inclusions (aggregates) undergo LLPS both *in vitro* and *in vivo*. Extensive research efforts have been put into understanding the relationship between the LLPS of proteins and their formation of toxic aggregates.^14,16,21,24^ These studies have revealed that the liquid condensates (droplets) mediate the conversion of soluble protein in the monomeric form to large pathological aggregates. Proteins undergo a liquid-to-solid phase transition within these droplets, due to high local protein concentration and reduced molecular motion as compared to the surrounding medium.^13^ The solid aggregates formed within the droplets act as nucleation sites for further protein aggregation and contribute to the process of neurodegeneration.

LLPS in proteins is primarily driven by weak intermolecular interactions promoting lower free energy of the system.^12,25^ IDPs possess a biased amino acid sequence, rich in polar, charged, and aromatic amino acids,^25–28^ which enable various intermolecular interactions including electrostatic, dipole-dipole, π-π, or cation-π interactions.^15,29–32^ These interactions play a crucial role in promoting the entropically unfavourable phase separation of IDPs. Also, IDPs exhibit high conformational flexibility, allowing them to attain the extended dynamic conformations associated with LLPS.^27,33^ In contrast, globular proteins typically fold into compact, three-dimensional structures stabilized by hydrogen bonding, hydrophobic, electrostatic, and other non-covalent interactions.^34^ The energy required to disrupt these intramolecular interactions and allow for intermolecular associations is usually high, making LLPS less likely. While folded proteins face these constraints, recent studies have reported that folded proteins can also undergo LLPS under specific conditions that alter the balance between intramolecular interactions and intermolecular associations.^35–37^ Motivated by these findings, our study aims to investigate the effect of crowding on the LLPS and aggregation behaviour of a globular protein, small ubiquitin-like modifier 1 (SUMO1).

SUMO1 is a 97-residue protein well-studied in terms of structure, stability, and function.^38–41^ It plays a crucial role in regulating various cellular processes through post-translational modification of its target proteins by SUMOylation.^42,43^ However, unlike ubiquitin, which is primarily associated with targeting proteins for <u>proteasomal degradation</u>, SUMO1 is involved in cellular processes such as nuclear transport, transcriptional regulation, apoptosis, and protein stability.^43,44^ The process of SUMOylation is carried out by a cascade of enzymes, which includes an E1 activating enzyme (SAE1/SAE2), an E2 conjugating enzyme (Ubc9), and an E3 ligase (such as PIAS or RanBP2). These enzymes recognize a specific sequence ΨKXD/E in target proteins, where Ψ is any hydrophobic residue, mostly isoleucine, leucine, or valine and X is any residue.^43^ The carboxylic group on the C-terminal of SUMO1 forms an iso-peptide bond with the ε-amino group of an acceptor target lysine residue. Interestingly, several IDPs involved in neurological diseases contain one or more SUMO1 recognition motifs and therefore can act as SUMO1 targets.^45,46^ Also, SUMO1 is found to be co-localized with IDPs in inclusion bodies isolated from patients’ brains suffering from neurodegenerative diseases such as multiple system atrophy, Huntington’s disease, and other related poly-glutamine disorders.^45–47^

In this study, we present the phase separation of SUMO1 under crowded conditions, at physiological pH, and room temperature. We find that SUMO1 undergoes conformational changes inside the phase-separated droplets which further mature to form solid aggregates. Our results imply that the relevance of LLPS extends beyond IDPs, indicating that folded proteins can also undergo LLPS under specific conditions.

## Experimental section: Methods

### 1. Expression and purification of proteins

SUMO1 variants (WT SUMO1, S1F66W, ΔN-SUMO1, ubiquitin (UBQ), SUMO2) were overexpressed in the BL21 (DE3) strain of *Escherichia coli* by inducing with 1mM IPTG (isopropylthio-D-1-thiogalactopyranoside) overnight at 25°C after the OD_600_ of the cell culture reached 0.6-0.8. The cells were harvested by centrifuging the cultures at 6000 rpm at 4°C for 30 minutes and the cell pellets were stored at −80°C. The cell pellets were resuspended in phosphate buffer (20 mM phosphate, 150 mM NaCl, pH 7.4) containing 0.1 mM PMSF (phenylmethanesulfonyl fluoride), triton-X, and other protease inhibitors such as leupeptin and pepstatin (∼0.1 mg both). The cells were lysed by sonication and centrifuged, and the supernatant was applied to a column of Ni-NTA-coated agarose beads (Qiagen). The beads were first washed with buffer and then with 20 mM imidazole (in buffer) pH 7.4. Finally, the proteins containing N-terminal His-tag were eluted with a buffer containing 300 mM imidazole at pH 7.4. The proteins were further purified using fast-performance liquid-phase chromatography (FPLC) Superdex 75 columns on Bio-Rad Biologic Duo-Flow FPLC system. The purified proteins were characterized by MALDI-TOF (Fig S1) and then stored at 4°C in the buffer.

### 2. Cysteine labelling with Alexa dyes

For labelling SUMO1 protein with Alexa488 dye, we prepared a 10 mM stock solution of dye in DMSO. The protein construct has three cystines (two terminal cysteines and one in the core of the protein at position 52). Alexa is expected to label any of these cysteines. Protein was incubated with 10X molar excess of TCEP for two hours. 10X molar excess of dye was then added to this protein solution containing TCEP and the reaction mixture was stirred for six hours in the dark at room temperature. After completion of the labelling reaction, the excess free dye was removed using a PD10 column. The labelled protein was then washed several times by a buffer exchange (with phosphate buffer) using a 3 kD MWCO Amicon membrane filter. For further experiments, we used a 1:100 molar ratio of unlabelled versus labelled protein.

### 3. Confocal transmission and fluorescence imaging

All imaging experiments were performed on a confocal laser scanning microscope, Zeiss Axio-Observer Z1 microscope (inverted) LSM 880. For the preparation of the phase-separated sample, we made a stock solution of 40% PEG-20 in buffer and adjusted its pH to 7.4. The sample was then prepared by adding the required volume of PEG-20 from the stock solution to the solution of SUMO1 (or S1F66W) such that the final concentration of PEG-20 and protein are 15% and 120-150 μM respectively. For transmission images, 50 μL of SUMO1 (or S1F66W) protein or the liquid droplets were placed onto a glass coverslip and images were acquired in Phase contrast mode using a 63x oil-immersion objective (numerical aperture 1.4). For fluorescence images, 1% of Alexa 488 labelled protein was added during the sample preparation and then confocal fluorescence images were taken at various time points using an excitation source of 488 nm. The emission light in the wavelength range of 500-620 nm was separated from the backscattered excitation light using a 488 long-pass dichroic mirror and collected using PMT. For ThT staining, a similar procedure was followed except that the sample was prepared using the unlabelled protein and a solution of ThT was added to it 15 min before imaging. All experiments were performed at room temperature in phosphate buffer (pH 7.4). Images were analysed and processed using ImageJ software (NIH, Bethesda, USA).

### 4. Fluorescence recovery after photobleaching (FRAP)

The sample preparation for FRAP experiments was done as described in Methods, section 3. FRAP studies were performed using a laser scanning confocal microscope (Zeiss Axio-Observer Z1 microscope) with a 63X oil immersion objective. Four different regions of interest (ROIs) were chosen: 1) The region to be photobleached in the middle of the droplet. 2) reference region in another droplet nearby to correct for passive bleaching during laser exposure. 3) a region outside the droplet to correct for background fluorescence intensity measurement. 4) a region containing all three ROIs for image acquisition before and after photobleaching. After acquiring five images at low-intensity laser settings, ROI-1 was photobleached at the 6th cycle using a 488 nm laser with 100% laser power and 50 iterations. Immediately after bleaching, the sample is monitored using the same low-intensity imaging settings as before the bleach. The recovery of fluorescence was recorded with an interval of 3s for 4-5 minutes until a plateau is reached. Time-dependent FRAP was performed by taking aliquots from the reaction mixture at mentioned time points.

For analysis, we used the method as described previously.^48,49^ The fluorescence values for every frame in the ROI-1 (F_bleach_) and ROI-2 (F_control_) were corrected for the background (F_bkg_) obtained from ROI-3.

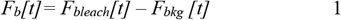

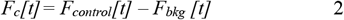

F_b_[t] is then corrected for passive bleaching to obtain experimental recovery R_norm_[t].

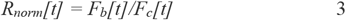

To obtain the fractional recovery, R_norm_[t] is then corrected by setting the first post-bleach value (t=0) to zero.

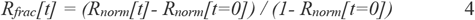

The data were then plotted and fitted using the single exponential recovery in Origin software.

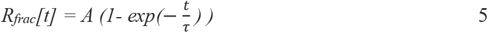

here, τ is the fluorescence recovery time constant, and ‘A’ corresponds to the mobile fraction of the fluorescent probe.

The half-time of the recovery (t_1/2_) was calculated from,

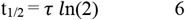

The mobile fraction (M.F) of the protein molecule in the ROI is depicted by;

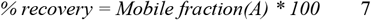

### 5. Transmission electron microscopy (TEM)

For TEM imaging experiments, the aggregated sample was diluted 20 times with water. 10 μL of the diluted sample was then directly adsorbed onto carbon-coated formvar grids (Electron Microscopy Science) for 5 min. We removed excess protein by placing EM grids two times on drops of Milli Q water. The samples were negatively stained by incubation in 2% (w/v) uranyl-acetate in water for 5 min, followed by brief washing in water and drying at room temperature overnight. The grids were subjected to transmission electron microscopy (TEM) using a 200 kV Tecnai-20 transmission electron microscope (TEM).

### 6. Steady-state fluorescence measurements

The steady-state fluorescence spectra of all the samples were recorded on a FluoroMax3 spectrofluorometer (HORIBA Jobin Yvon). For the preparation of aggregates, 120-150 μM of SUMO1 protein was incubated with 15% PEG-20 for more than six hours. For measuring tryptophan fluorescence, samples were excited at 295 nm and the emission spectra were recorded from 315 to 450 nm. For comparison, fluorescence spectra at t=0, immediately after the addition of PEG-20 was also measured. Measurement of ThT (final concentration of ThT in a sample was 30 μM) fluorescence was done by exciting the samples at 440 nm and recording the emission spectra from 460 to 600 nm. The excitation and emission bandwidths were kept constant within 2 to 3 nm for both (soluble and aggregated) samples. The excitation wavelength used for ANS fluorescence measurements was 385 nm and emission was collected from 400-700 nm. For all the measurements, the scan speed was 1 nm/s and the emission spectrum was averaged for three measurements at identical conditions. All experiments were done at room temperature.

### 7. Time-dependent scattering/fluorescence measurements

Kinetic fluorescence measurements were performed using a co-star black-coated 96-well plate on TECAN Infinite 200 Pro plate reader at 25 °C. The LLPS-mediated aggregate formation was initiated by adding 15% PEG-20 to 120-150 μM of protein sample. A sample volume of 200 μL was used for these measurements. The excitation and emission slits were kept at 9 and 20 nm, respectively. For tryptophan fluorescence in S1F66W, the excitation wavelength was 295 nm and emissions were collected at 337 nm (gain-90) every 1 min. For time-dependent ThT fluorescence measurements, the concentration of ThT in samples was kept at 8-10 μM. The excitation wavelength was 440 nm and emission (gain-110) was collected at 485 nm after every min. ANS fluorescence measurements with time were done at excitation wavelength 385 nm and emission was recorded at 500 nm, the concentration of ANS in a sample was 20 μM. The measurement is started immediately after the addition of PEG-20 (t=0) and is measured until the sigmoidal curves reach their saturation value (around 8-10 hours). The kinetic traces were plotted using Origin 2018. The droplet formation was monitored by recording the absorbance at 600 nm, at 25 °C on a Multiskan Go (Thermo Scientific) plate reader using 96-well polystyrene plates. Measurements were made after an interval of 1 min for 4-6 h until it reaches a saturation value. The data is normalized without any blank correction.

### 8. Circular dichroism measurements

The steady-state far-UV CD data were recorded on a Jasco J-1500 spectrometer. For sample preparation, 90 μM of SUMO1 protein was incubated with 15% PEG. After 2h and 6h of incubation, a small volume of sample was diluted six times with phosphate buffer and spectra were recorded in the range of 200-260 nm, with a 1 mm path length cuvette. For comparison with native SUMO1, spectra in the same range were recorded for 15 μM of SUMO1 protein in phosphate buffer.

### 9. Time-resolved fluorescence measurements

The time-resolved fluorescence of S1F66W was determined using the time-correlated single-photon counting (TCSPC) technique as described previously.^50^ The TCSPC data were recorded using a home-built setup that utilizes a pico-second dye laser. Samples were excited at 295 nm, and emission was collected at 337 nm. Fluorescence decay data were collected such that the peak count was >10,000. Fluorescence decays were fitted to three exponentials by minimizing χ^2^. Average fluorescence lifetime (*τ*_avg_) was estimated using the expression, 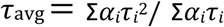, where *α*_i_ and *τ*_i_ are amplitude and lifetime of the ith component in the decay analysis.

### 10. Line-confocal Raman measurements

Raman measurements were performed using a home-built line-confocal Raman microscope as described previously.^51^ In brief, the microscope uses a commercially sourced laser (Verdi V-10) which gives up to 10 W output power of 532 nm wavelength, a Horiba iHR320 spectrometer and a fluorescence microscope (Olympus IX71). The beam waist of the laser source coming from the Verdi-V10A is increased using a telescope setup consisting of a pair of convex lenses (L_1_: f=5 cm, L_2_: f=20 cm, f= focal length of the lens). A cylindrical lens (CL) was used to create the line-shaped beam. The beam was focussed using a 1.2 NA 60X objective (Olympus) onto the sample. A 532-long pass dichroic mirror (DM, LP03-532RE-25, Semrock) was used to separate the excitation photons and stokes-shifted Raman photons. The Raman photons were focused at the entrance of the spectrometer using a Tube lens of 16 cm focal length. Inside the spectrometer, the photons of different wavelengths were separated by a grating and imaged by a synapse Charge Coupled Device (CCD) camera. The wavenumbers were calibrated using sulphur powder and naphthalene as a calibrant. For measuring Raman spectra of SUMO1 samples (native state and aggregated state), glass coverslips of thickness 13-20 mm containing the sample were kept on top of the water-immersed objective. The line averaged Raman spectra were acquired for 1 h with 200 mW excitation laser power at the back aperture of the objective.

## Results

### SUMO1 undergoes LLPS under crowded conditions

To investigate the behaviour of globular protein SUMO1 (Fig 1A) in crowded conditions, we subject it to the inert crowding agent PEG (Molecular weight 20kD unless specified). All the experiments are performed at room temperature and in phosphate buffer (pH 7.4). We observe visual turbidity in the solution of SUMO1 protein within a few minutes after incubation with PEG (Fig 1B). The optical transmission image of this turbid solution shows the presence of small droplets of ∼1-2 μm size, suggesting liquid-liquid phase separation (LLPS) of the protein. We then labelled SUMO1 with a fluorescent dye (Alexa488) to confirm the presence of SUMO1 in the droplets. Labelled and unlabelled proteins are mixed in a 1:100 molar ratio and confocal images are acquired at different time points after adding PEG (Fig 1C). Initially, we observe fluorescent droplets of spherical shape (t = 10 min) which then become irregular larger droplets (t = 2 hr) and finally form bigger aggregates (t = 5 hr). To further confirm the LLPS, we perform FRAP (Fluorescence Recovery After Photobleaching) experiments at different time points and then probe the changes in dynamics of SUMO1 with the maturation of droplets (Fig 1D). The raw data obtained in the FRAP experiment is then analysed as described in Methods (section 4) and the fluorescence recovery curves shown in Fig 1D are then used to calculate % recovery (Fig 1E) and half-time of recovery (t_1/2_) (Fig 1F). At t = 15 min, we observe rapid fluorescence recovery of 80%, in agreement with their liquid-like nature. However, we observe a decrease in % recovery and an increase in t_1/2_ with the ageing of droplets. This is expected due to the decreased molecular diffusion of SUMO1 inside the droplets owing to its aggregation. We also use other crowding agents like PEG-8kD and Dextran-20kD and added them to a solution of SUMO1. The appearance of fluorescent droplets verifies that the LLPS effect of SUMO1 is not specific to PEG-20 (Fig 1G). Taken together, these results demonstrate that SUMO1 protein undergoes condensation into liquid-like droplets in the presence of crowding agents.

**Fig 1.**
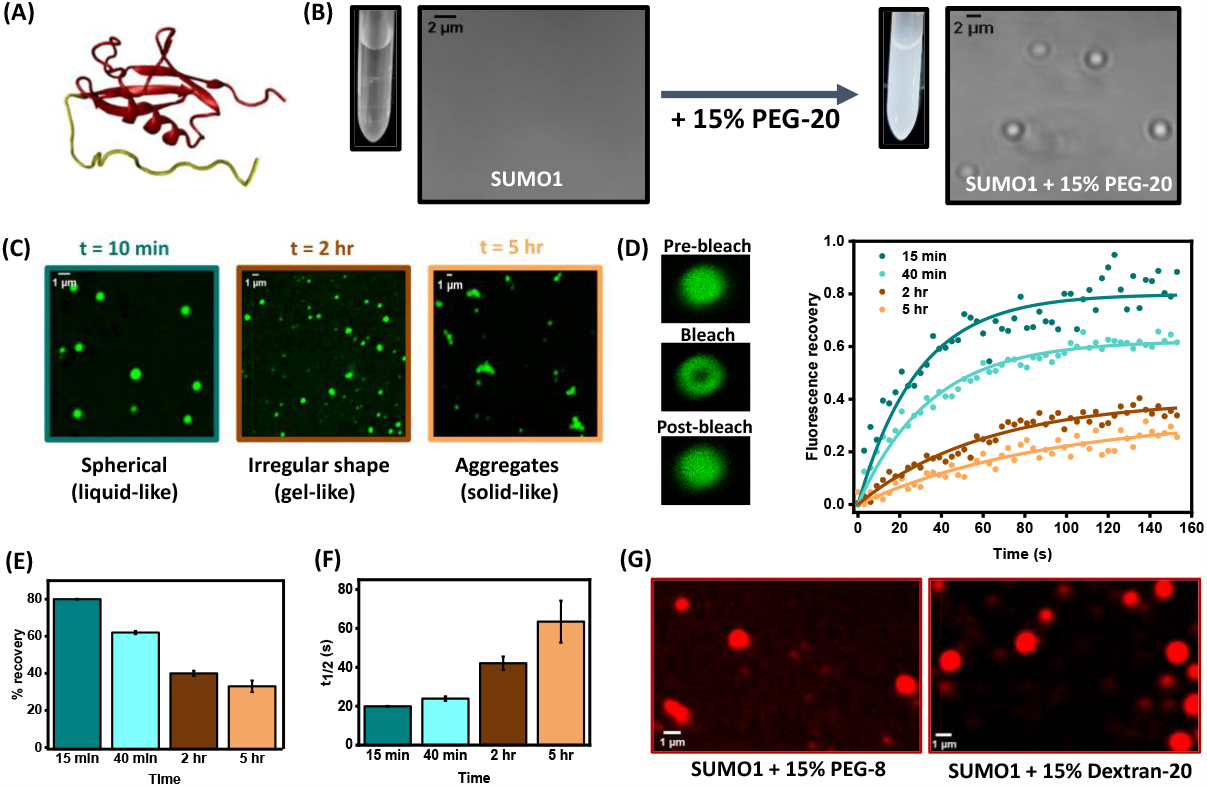
SUMO1 undergoes LLPS under crowded conditions. (A) NMR structure of SUMO1 (PDB ID: 2N1V) showing the structural domain in red and the N-terminal tail in yellow. (B) Optical transmission images of SUMO1 (120 μM) in the absence (left) and presence (right) of PEG-20kD (15%), within 1 hr after addition. (C) Confocal fluorescence images of 120 μM Alexa488-labelled SUMO1 at different time points after incubation with PEG-20kD (15%). FRAP measurements of SUMO1 droplets at indicated times (time after addition of PEG) to measure the change in dynamics of droplets (left panel: FRAP protocol). (E, F) % recovery and t_1/2_ at different time points, calculated from FRAP recovery curves. (G) Confocal fluorescence images of 120 μM Alexa488-labelled SUMO1 with PEG-8kD (left) and Dextran-20kD (right). All the experiments were performed in phosphate buffer (pH 7.4), 150 mM NaCl, at room temperature.

### Phase transition of liquid condensates forming amorphous aggregates

We measure Thioflavin-T (ThT) fluorescence in the absence and presence of SUMO1 aggregates (Fig 2A). ThT is reported to show enhanced fluorescence on binding to amyloid-like aggregates. The SUMO1 aggregates show enhanced ThT fluorescence suggesting an amyloid-like structure. We then compare the kinetics of the formation of aggregates with that of droplet formation (Fig 2B). The formation of droplets is monitored by standard kinetic assays based on turbidity (scattering) measurements at 600 nm and the formation of aggregates is monitored by a change of ThT fluorescence at 485 nm. The slower kinetics of change in ThT fluorescence as compared to changes in turbidity of solution supports the fact that droplet formation precedes the aggregation of SUMO1. Also, confocal images of the ThT-stained sample of SUMO1 condensates verify that the formation of aggregates initiates inside the droplets (Fig S2). We also measure changes in ANS fluorescence with time after incubation of SUMO1 protein in PEG (Fig S3). We do not observe any significant changes in fluorescence intensity as the protein aggregates. This is probably due to the preferential binding of ANS to PEG present in the sample solution. ANS fluorescence remains quenched in the buffer but increases in the presence of 15% PEG with a blue shift from 524 to 506 nm (Fig S3). Furthermore, to gain insights into the structural properties of SUMO1 in aggregates, we recorded Raman spectrum of SUMO1 aggregates and compared it with SUMO1 in solution (Fig 2C, 2D). Both spectra are normalized to the phenylalanine peak at 1003 cm^-1^. SUMO1 in its native state exhibits an amide band at 1654 cm^-1^ which shifts to 1665 cm^-1^ upon aggregation. This suggests an alpha-helix to beta-sheet transition in the aggregated state of the SUMO1 protein. Furthermore, we observe a decrease in the spectral broadening of the amide-I band at 1654/1665 cm^-1^ in aggregates, indicating that there is less conformational heterogeneity in SUMO1 aggregates compared to the native state. While ThT fluorescence and Raman studies indicate the presence of amyloid-type aggregates, morphology from TEM images (Fig 2E, 2F) suggests the aggregates to be amorphous. This implies that aggregates are rich in beta-sheet secondary structures but are of amorphous morphology.

**Fig 2.**
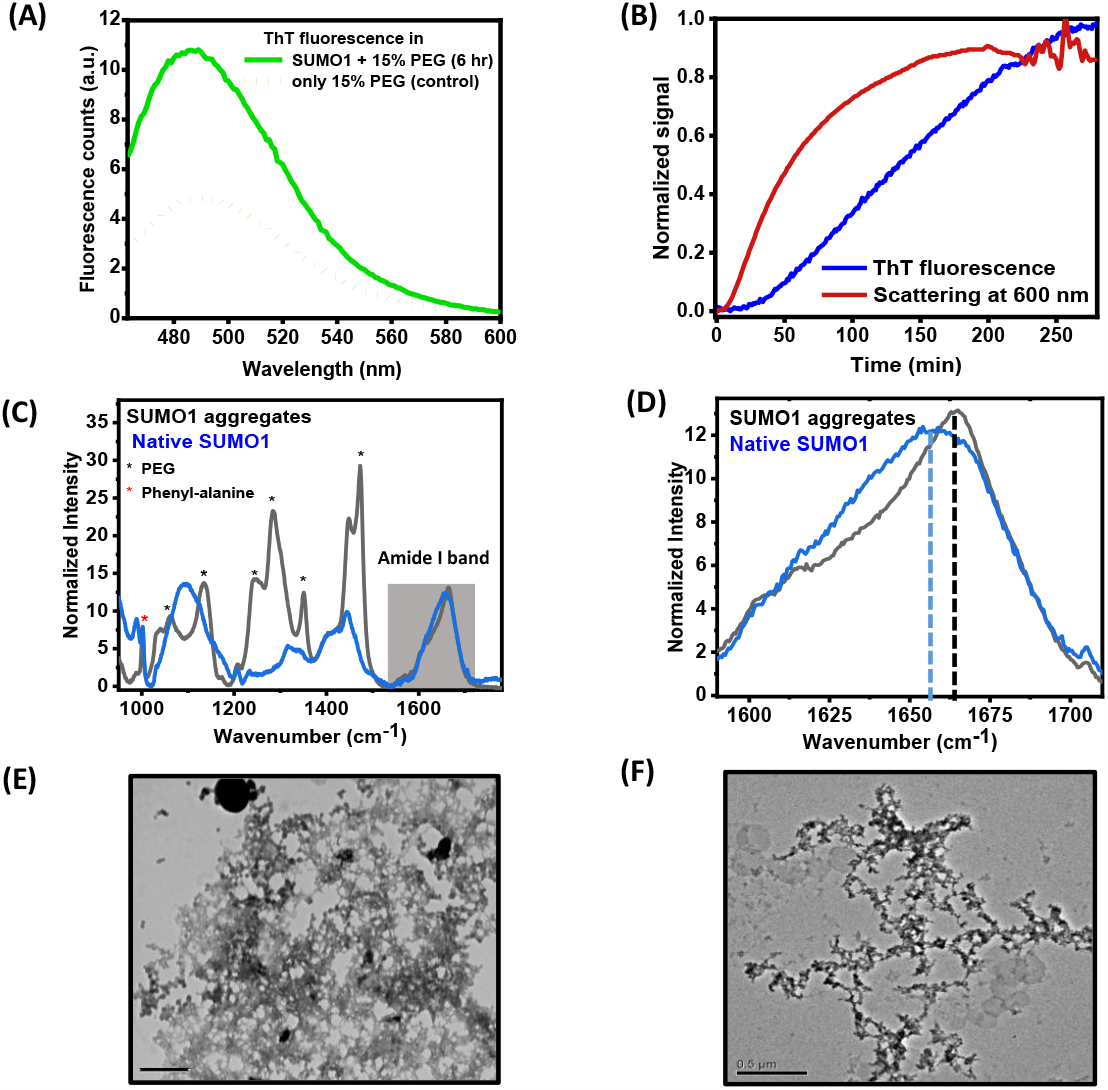
Phase transition of liquid condensates of SUMO1 forming amorphous aggregates. (A) Fluorescence spectra of Thioflavin-T (ThT) in 15% PEG-20kD (dashed green line) and ThT-incubated with SUMO1 aggregates (solid green line), excitation wavelength used is 440 nm. (B) ThT fluorescence (blue) intensity at 485 nm (excitation wavelength -440 nm) and scattering (red) measurements at 600 nm with time after incubating 120 μM SUMO1 with PEG-20kD (15%), the concentration of ThT in a sample is 10 μM (C) Raman spectra of native SUMO1 (blue) and SUMO1 aggregates (black) normalized w.r.t phenylalanine peak (red asterisk) at 1003 cm^-1^. (D) zoomed-in region of the amide-I band from (C). (E, F) Transmission electron micrographs of negatively stained SUMO1 aggregates formed at two different conditions, after incubating 120 μM SUMO1 with 15% PEG-20kD for 8 hours, scale bar is 100 nm (E) and 50 μM with 15% PEG-20kD for 11 days, scale bar is 500 nm (F).

### Conformational changes of SUMO1 within condensates promote aggregation

To probe the conformation changes associated with the phase separation of SUMO1, we synthesize its tryptophan mutant S1F66W where phenylalanine at the 66^th^ position is mutated with tryptophan. The fluorescence intensity of a fluorophore is sensitive to its environment, which suggests that changes observed in the tryptophan fluorescence intensity can be correlated to the protein conformational changes. It has already been reported by our group that this mutant is similar to wild-type (WT) SUMO1 in terms of its secondary and tertiary structure along with its stability.^52^ Tryptophan fluorescence of S1F66W remains quenched in its native state and increases on unfolding with red-shift (around 20 nm) in its emission maximum.

S1F66W also shows similar phase separation behaviour to WT SUMO1 (Fig S4) on the addition of PEG. We record tryptophan fluorescence spectra of S1F66W before and 6 hr after incubation with PEG (Fig 3A). We observe an increase in fluorescence intensity with very little (∼3 nm) red shift in emission maximum. The increase in the fluorescence intensity suggests a rearrangement of surrounding residues around the Trp. On the other hand, very little change in the emission maximum suggests a similar polar environment or solvent exposure around the Trp. This observation is substantiated by time-resolved measurements (Fig 3B) where the average fluorescence lifetime also increases from 0.54 to 1.57 ns. (Supplementary table S1) in the presence of PEG, which is in correlation with steady-state fluorescence measurements. The considerable increase in fluorescence intensity and lifetimes of Trp in aggregated protein supports the fact S1F66W undergoes structural changes in the presence of PEG and it attains some non-native conformation. Furthermore, the supernatant obtained by centrifugation of the aggregated solution of S1F66W exhibited a similar fluorescence decay curve as of native S1F66W, suggesting that the soluble protein fraction has a native-like structure. We further monitor the kinetics of Trp fluorescence at 337 nm and compare the rate with other probes (scattering and ThT fluorescence) (Fig 3C). We find that the conformational changes (Trp fluorescence) occur at a faster rate than the aggregation (ThT), but slower than the droplet formation (scattering at 600 nm).

**Fig 3.**
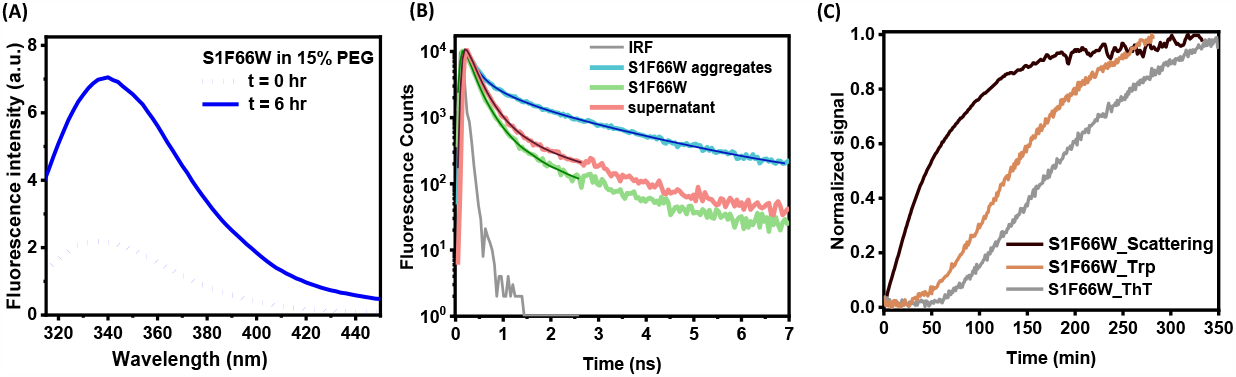
Conformational changes of SUMO1 within condensates promote aggregation. These experiments are performed with S1F66W, a tryptophan mutant of SUMO1. (A) Tryptophan fluorescence spectra of S1F66W at t=0 (dashed blue line) and 6 hr (solid blue line) after incubation with PEG-20kD, excitation wavelength used is 295 nm. (B) Fluorescence decay curves of native S1F66W (green), S1F66W aggregates formed after incubation with 15% PEG-20kD for 6 hr (blue), and the supernatant obtained after centrifuging the aggregated S1F66W solution (pink). (C) Thioflavin-T fluorescence intensity (grey), tryptophan fluorescence (orange) and scattering (black) measurements with time after incubating 150 μM S1F66W with PEG-20kD (15%).

### LLPS of SUMO1 is reversible

We measure CD spectra of droplets (2 hr incubation with PEG) and aggregates (6 hr incubation with PEG) of SUMO1 at different time points after diluting with buffer (Fig 4A, 4B). In both cases, the CD spectra immediately after dilution show less signal compared to native SUMO1 of the same concentration, however, the spectral features were similar. It might be due to the signal being dominated by the soluble fraction of protein, with negligible contribution from the droplet or aggregate phase. Therefore, we could not get information about the secondary structure of a protein in the droplet or aggregated phase. Interestingly, the initial lower signal of droplets compared to the native SUMO1 protein increased with time after dilution and overlapped with that of native SUMO1 in ∼3 min. This suggests that the LLPS is reversible and droplets dissolve back in the solution increasing the amount of soluble protein and thus increase the signal. However, we did not observe such an increase in signal with time for a diluted sample of aggregates, indicating that the aggregate formation is irreversible in the time scale of 30 min.

**Fig 4.**
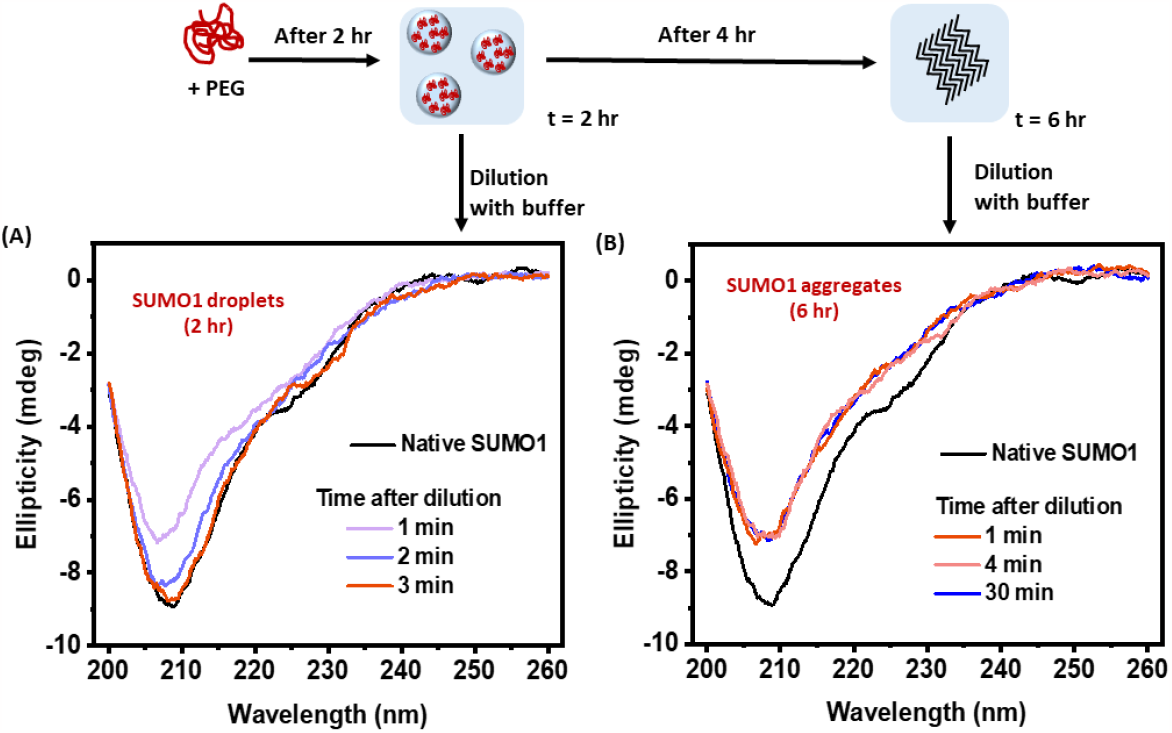
LLPS of SUMO1 is reversible. (A) CD spectra measured with time after dilution of SUMO1 droplets (2 hr incubation with PEG-20kD) and its comparison with CD spectra of native SUMO1 (in black, same concentration as of diluted samples). (B) CD spectra measured with time after dilution of SUMO1 aggregates (6 hr incubation with PEG-20kD) and its comparison with CD spectra of native SUMO1 (in black, same concentration as of diluted samples).

### The disordered tail of SUMO1 is not essential for phase separation/transition

SUMO1 has a long unstructured N-terminal tail (Fig 1A). To investigate the effect of this N-terminal tail on phase separation and aggregation properties, we used a SUMO1 variant (ΔN-SUMO1, 17 residues deleted from the N-terminus) as shown in Fig 5A. ΔN-SUMO1 has been earlier shown to have similar thermodynamic stability as full-length SUMO1.^52,53^

**Fig 5.**
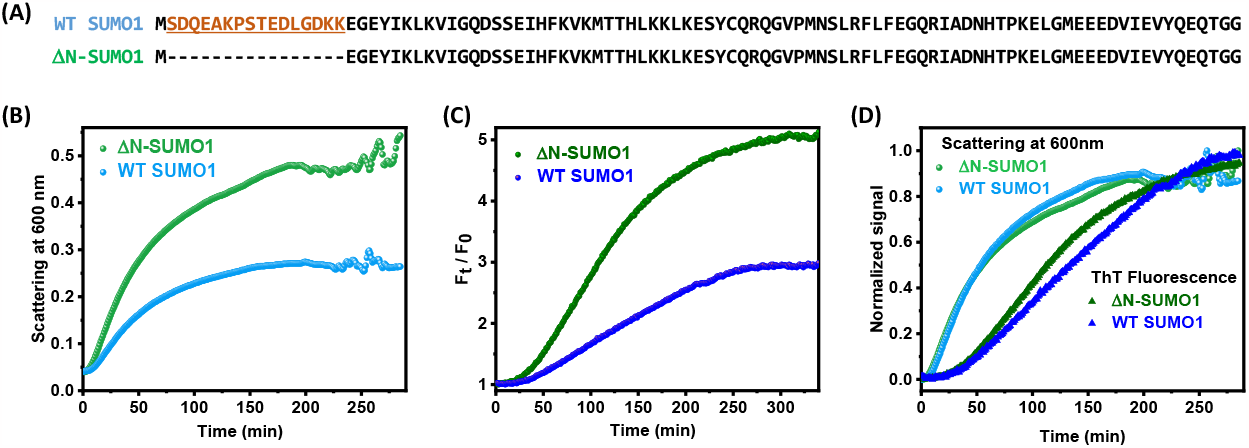
The disordered tail of SUMO1 is not essential for phase separation/transition. (A) Sequences of WT SUMO1 and ΔN-SUMO1. (B) Scattering measurements with time after incubating 120 μM SUMO1 (blue) and ΔN-SUMO1 (green) with PEG-20kD (15%). (C) Thioflavin-T fluorescence intensity measurements with time after incubating 120 μM SUMO1 (blue) and ΔN-SUMO1(green) with PEG-20kD (15%). (D) Normalized plots of 5B and 5C.

We compare ΔN-SUMO1 and full-length SUMO1 in terms of their tendency to phase separate and aggregate under similar crowded conditions. For this purpose, we incubate both proteins with PEG and all other parameters such as concentration of PEG and the proteins, pH, and temperature were kept constant. We monitor the time course of SUMO1 and ΔN-SUMO1 phase separation and aggregation by turbidity and ThT fluorescence measurements. Interestingly, we observe that the overall rate of phase separation (Fig 5B) and aggregation (Fig 5C) is more in the case of ΔN-SUMO1 compared to the full-length SUMO1. However, the rate constants, as evident from the normalized plots, were very similar (Fig 5D). Also, the maximum value of F_t_/F_0_ and scattering (Fig 5B, 5C) is more (almost twice) in the case of ΔN-SUMO1 indicating the presence of more aggregates compared to full-length SUMO1. We then centrifuge the aggregated solutions of full-length SUMO1 and ΔN-SUMO1 to determine the concentration of residual protein (final concentration) in the supernatant (Fig S5). Less concentration of residual protein in the case of ΔN-SUMO1 compared to full-length SUMO1 supports that more amount of the protein aggregates in the case of ΔN-SUMO1 for the same initial concentrations of protein and PEG. Overall, these results highlight that the truncated variant of SUMO1, lacking any IDR also undergoes LLPS and aggregation.

## Discussion

SUMO1 is a structured globular protein which is highly soluble in dilute buffer conditions. In this work, we show that SUMO1 undergoes rapid LLPS under crowded conditions at room temperature and physiological pH (Fig 1). The observed effect of crowding-induced phase separation of SUMO1 may be due to either of the two reasons: first, increased intermolecular interaction in SUMO1 due to volume exclusion by PEG polymer and second, specific interaction between SUMO1 and PEG molecules leading to their co-condensation. Similar effects are also observed in another crowding agent Dextran (Fig 1G), suggesting that the effect is mostly due to volume-exclusion. Crowding agents are added in almost all *in vitro* phase separation studies. In our study of SUMO1, we do not observe phase separation in the absence of PEG even at very high concentrations (400 μM) of the protein. In many previous studies, it is observed that LLPS occurs even in the absence of crowders but the addition of crowders lowers the critical concentration required for phase separation. As an example, the critical concentration for LLPS of alpha-synuclein lowered from 500 μM to 200 μM in the presence of 10% PEG^14^. In contrast, crowding environment is essential for SUMO1 to undergo LLPS. Similar observations were reported for ptau441, where no LLPS is observed without the addition of PEG.^23^ We further verified the liquid-like behaviour of the phase-separated condensates of SUMO1 by their fast fluorescence recovery in FRAP experiments (Fig 1D). Within a few minutes, however, the condensates become gel-like and then finally solid aggregates are formed which show almost no fluorescence recovery after photobleaching. These solid SUMO1 aggregates show enhanced ThT fluorescence and increased β-sheet content (Fig 2A, 2D) but morphology obtained from TEM images (Fig 2E, 2F) suggests that the aggregates are amorphous.^54^ Using ThT fluorescence as a probe, we measured the kinetics of aggregation of SUMO1 at various concentrations of PEG (Fig S6). We find that the extent of SUMO1 aggregation and lag phase is dependent on PEG concentration.

Aggregation of SUMO1 in the presence of PEG is associated with conformational changes as evident from the shift in the amide-I band in Raman spectra (Fig 2C, 2D) and the increase in tryptophan fluorescence (Fig 3A). By comparing the kinetics of all steps obtained from various spectroscopic probes (Fig 3C), we have proposed a model for SUMO1 phase separation and aggregation (Fig 6). Since the rate of increase in tryptophan fluorescence with time is less than that obtained from the scattering measurements (Fig 3C), we deduce that conformational changes occur after the formation of droplets. Also, the protein in the dilute phase retains its native conformation as confirmed by excited-state lifetime values of tryptophan obtained from the supernatant (Fig 3B). This suggests that the environment in the vicinity of the protein in the droplets (dense phase) is different than that at the outside of them (dilute phase). Therefore, the protein undergoes a conformational change to minimize its free energy inside the droplets. From time-dependent ThT fluorescence measurements, we observe that the kinetics of aggregation is slower than that of conformation change reported by the tryptophan. This suggests that the SUMO1 molecules in non-native conformation inside the droplets interact with other molecules and this self-association leads to the formation of solid aggregates. The very fast reversibility of the droplet formation in the initial stages suggests that the intermolecular interactions among the proteins in the droplet are weak (Fig 4A). On the other hand, the aggregation (after 6 hr) is irreversible, indicating changes in the structure leading to a strong intermolecular association (Fig. 4B).

**Fig 6.**
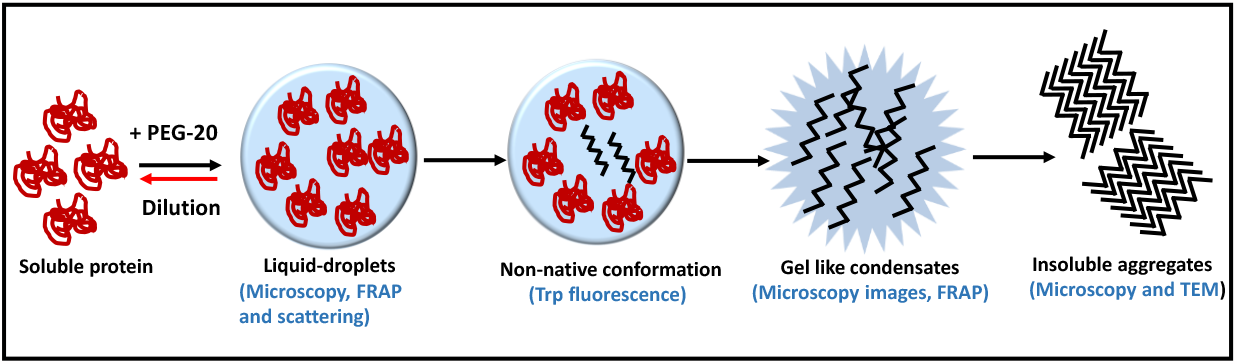
Schematic of SUMO1 phase separation and aggregation in the presence of PEG illustrating the different stages from soluble protein to aggregation via LLPS. Formation of droplets and further conformational changes inside the droplets leading to aggregation. LLPS is in equilibrium with soluble protein, whereas aggregate formation is irreversible.

It must be noted SUMO1 has an N-terminal flexible disordered tail. As already mentioned, LLPS is commonly associated with proteins that have IDRs or proteins with low-complexity sequences and thus it is important to investigate if this N-terminal tail drives phase separation and aggregation of SUMO1 protein. For that purpose, we have used a deletion mutant of SUMO1 lacking the unstructured region (Fig 5A). Similar rate constants obtained for both ΔN-SUMO1 and full-length SUMO1 (Fig 5D) from ThT fluorescence and turbidity measurements suggest that the intrinsic tendency of the protein to phase-separate an aggregate does not change after the removal of the disordered segment. Interestingly, under similar concentrations of PEG and protein, a higher fraction of the protein partitions into the dense phase for ΔN-SUMO1 compared to the full-length protein (Fig 5B, 5C and S5). This resulted in the formation of more aggregates and less residual protein in the supernatant. Earlier, Grana-Montes and coworkers demonstrated the role of the N-terminal flexible tail of SUMO proteins in their aggregation to amyloid structures at a high temperature.^53^ Using rate constants obtained from ThT and CD studies, they also showed that the removal of this N-terminal tail increases the aggregation propensity of SUMO proteins. In their study, however, rate constants were also more for the deletion mutant which suggests that the mechanism of aggregation is different in crowded conditions compared to destabilizing conditions. Also, the phase separation of ΔN-SUMO1 highlights the fact that the presence of IDRs is not an essential requirement for phase separation. Although it is well established that IDRs provide more conformational flexibility to proteins to develop weak associative interactions required for phase separation, our study shows that globular proteins lacking any IDRs can also phase separate by inter-molecular interactions induced by crowding.

Other ubiquitin-family proteins, for example, SUMO2 and ubiquitin have similar structures as SUMO1, however their amino-acid sequences are different. We compared their LLPS propensity with SUMO1 to understand if the effect is specific to the SUMO1 sequence (Fig S7). At the same concentrations of protein and PEG, ubiquitin did not phase-separate in the timescales required for SUMO1 phase-separation or aggregation. For SUMO2, some increase in the scattering was observed but ThT fluorescence did not show any change. This does not, however, exclude the possibility of these proteins to phase separate under crowded conditions. It might be possible that they have less propensity for LLPS and might phase separate at higher protein/ PEG concentrations or at longer time scales. Also, this suggests that the sequence of amino acids in SUMO1 makes it uniquely more prone to LLPS and aggregation under crowded conditions, compared to other proteins of the same family with similar structures. To determine if denatured SUMO1 can undergo LLPS, we mixed SUMO1 with denaturants like urea and GdHCl and then incubated it with PEG-20 (Fig S8). The denatured protein does not show LLPS that implies crowding-induced LLPS effects require a native-like structure of SUMO1 protein.

Patel et al. previously demonstrated phase separation of HSA, a globular protein, in the presence of PEG.^36^ They show that denatured HSA does not undergo phase separation highlighting the importance of the native protein in intermolecular interactions. However, the liquid droplets of HSA do not undergo a transition to form solid aggregates even up to 60 days at lower (<500 μM) protein concentrations which contrasts with our as well as other studies on LLPS of proteins. Erika Bullier-Marchandin and co-authors recently reported LLPS and aggregation of BSA in the presence of PEG.^37^ They found no changes in the amide-I band (by FTIR) during protein phase separation and aggregation, unlike our observations of increased beta-sheet content in SUMO1 aggregates. Here, we demonstrated that the LLPS of SUMO1 is not specific to PEG and occurs with other crowders such as Dextran, which was not addressed in previous studies. This is very crucial because while PEG is typically considered an inert crowder, it can interact with the protein’s hydrophobic residues in certain cases.^55^ While the above-mentioned studies report the LLPS of HSA and BSA, their biological relevance is not immediately apparent given that these are serum proteins. On the other hand, our results showing LLPS and aggregation of SUMO1 may have significance in neurodegenerative diseases. Many IDPs involved in various neurodegenerative diseases are SUMO substrates and SUMO proteins have been isolated from inclusion bodies from such patients. ^45–47,56^ A few examples are alpha-synuclein (in PD), Tau protein (in AD), Huntington (in Huntington’s disease) and amyloid precursor protein APP (in AD). Wu, J. *et al*. showed that amorphous aggregates from other proteins (HAS, BLG, SOD1) may interact with Aβ and lead to inhibition of fibrillation in favour of non-fibrillar aggregation.^57^ Furthermore, it has been shown that aggregation of IDPs such as alpha-synuclein and tau protein into amyloid-like structures occurs through liquid-liquid phase separation which is observed to take place *in vitro* in a few days under crowded conditions. In contrast, SUMO1 phase separates within a few minutes in the presence of PEG. We hypothesize that the presence of the SUMO1 tag may be of great importance in modulating the propensity of phase separation or aggregation of its IDP substrates under diseased conditions.

## Conclusion

In our study, we successfully demonstrated the rapid phase separation of SUMO1 protein under crowded conditions, highlighting the importance of its structured part in driving LLPS. It is believed that intrinsically disordered regions (IDRs) are the sole drivers of LLPS, while the structured part alone cannot induce phase-separation. However, our results suggest that structured proteins can overcome their biophysical constraints and undergo phase separation under specific conditions. Our findings have the potential to alter the existing prospective and encourage further investigation into the liquid-liquid phase separation (LLPS) of structured proteins to gain deeper insights into the key drivers of the process. Also, LLPS of SUMO1 may be of great biological relevance in the context of neurodegenerative diseases and future studies on the effect of SUMO1 tag on LLPS of IDPs, that possess SUMOylation sites, may improve our understanding of the disease mechanisms.

## SUPPORTING INFORMATION

Supporting information contains an additional data/figures.

## ACKNOWLEDGEMENTS

S. R. K. A. acknowledges the financial support provided by the Department of Atomic Energy (DAE), India, through Project No. 12-R&DTFR-5.10-0100. The authors thank Rudheer D Bapat, Shilpa Shirodkar, Jayesh Parmar, and Lalit Borde for assisting with TEM imaging. We also thank the assistance of Geetanjali A. Dhotre in recording the mass spectra using the matrix-assisted laser desorption/ionization–time-of-flight (MALDI-TOF) instrument, and the support of Mamata V. Kallianpur in conducting TCSPC fluorescence lifetime experiments. We are grateful to Debanjana Das and Suman Tiwari for their valuable comments and suggestions.

## For Table of Contents Only (TOC)

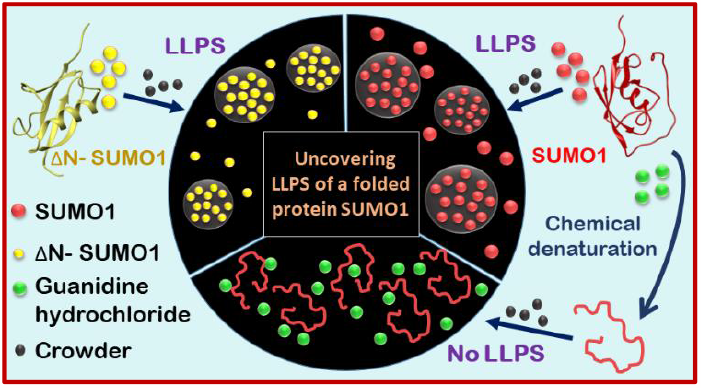

