## Supporting Information for "Liquid-liquid phase separation and aggregation of a globular folded protein SUMO1"

**Author ORCIDs**

Simran Arora: 0000-0002-6423-4442

Sri Rama Koti Ainavarapu: 0000-0002-1646-2731

### Supplementary Figures

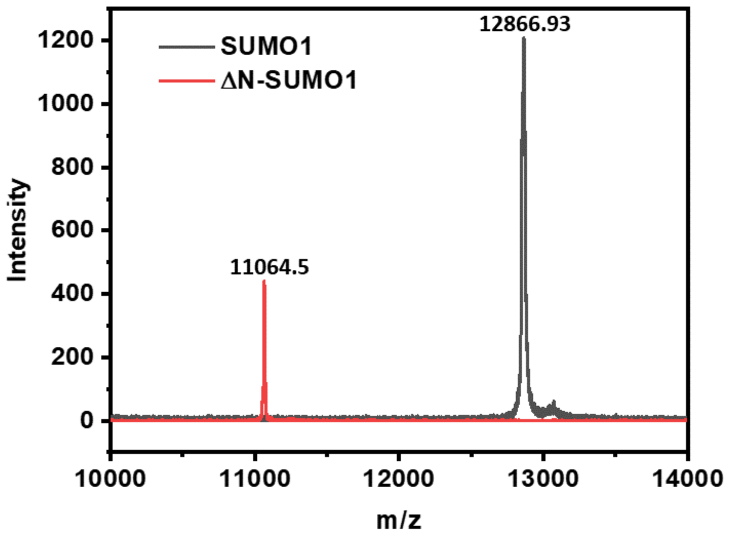


SUMO1- MRGSHHHHHHGSSDQEAKPSTEDLGDKKEGEYIKLKVIGQDSSEIHFKVKMTTHLKKLKESYCQRQGVPMNSLRFLWEGQRIADNHTPKELGMEEEDVIEVYQEQTGGRSCC

ΔN-SUMO1- MRGSHHHHHHGS -------------------------- EGEYIKLKVIGQDSSEIHFKVKMTTHLKKLKESYCQRQGVPMNSLRFLWEGQ RIADNHTPKELGMEEEDVIEVYQEQTGGRSCC

**Fig S1**. MALDI-TOF of SUMO1 and ΔN-SUMO1 after purification through Ni-NTA affinity column and SEC column. Full sequence of SUMO1 and ΔN-SUMO1 used to express the proteins using $His_6$-tag. Additional residues are from restriction sites used in pQE80L vector.

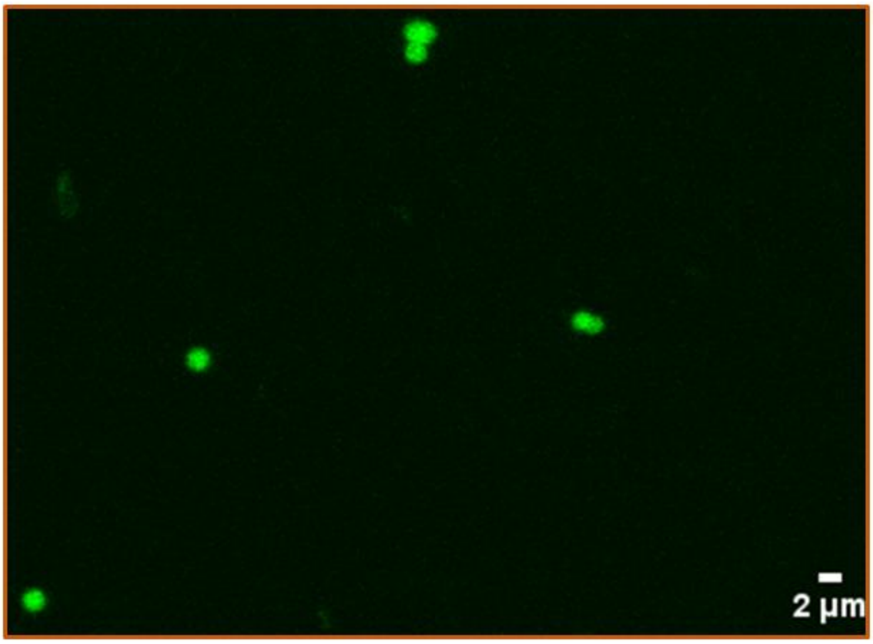


**Fig S2**. Confocal fluorescence images of ThT-stained SUMO1 sample after 1 hr incubation with PEG (15%).

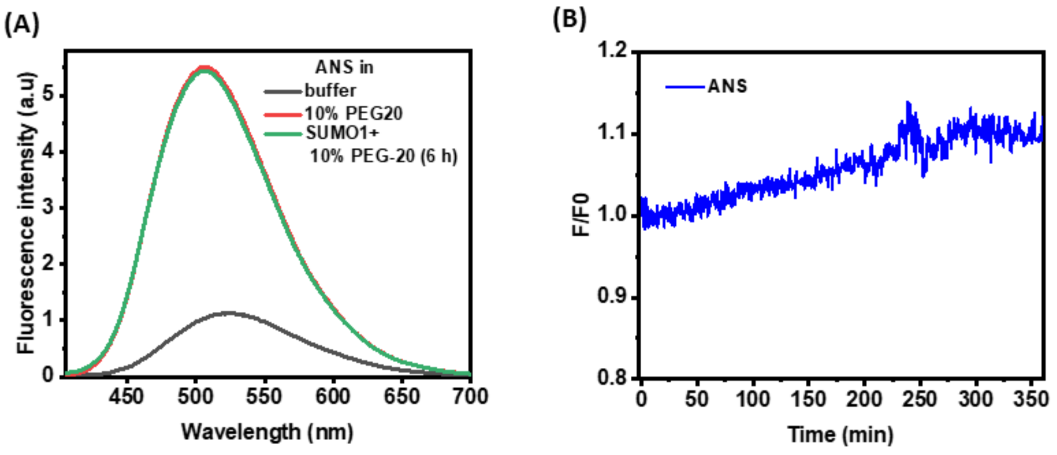


**Fig S3**. (A) ANS fluorescence spectra with and without SUMO1 protein in 10% PEG-20. (B) ANS fluorescence measured (at 500 nm) with time after incubation of 120 μM with 10% PEG-20. Excitation wavelength used is 385 nm.

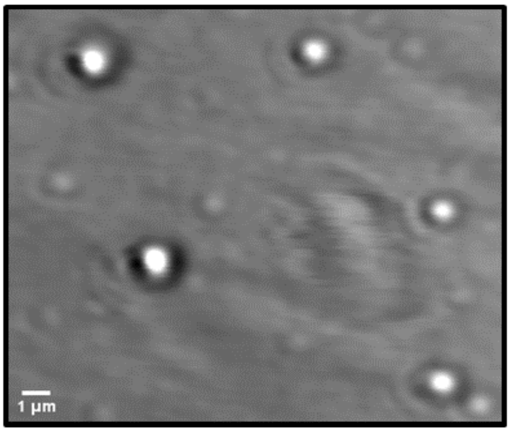


**Fig S4.** Optical transmission image of S1F66W droplets formed after addition of 15% PEG-20 to 120 μM of the protein.

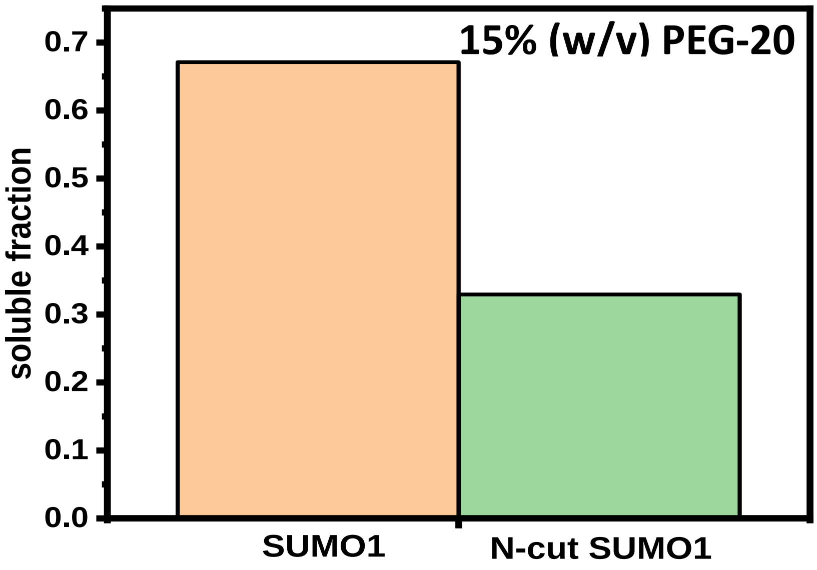


**Fig S5**. Soluble fraction (final/initial) obtained in the supernatant after incubation of 125 μM of protein with 15% PEG overnight. Concentrations were calculated from absorbance measurements using $\varepsilon_{295}$= 4470 $M^{-1}$ $cm^{-1}$.

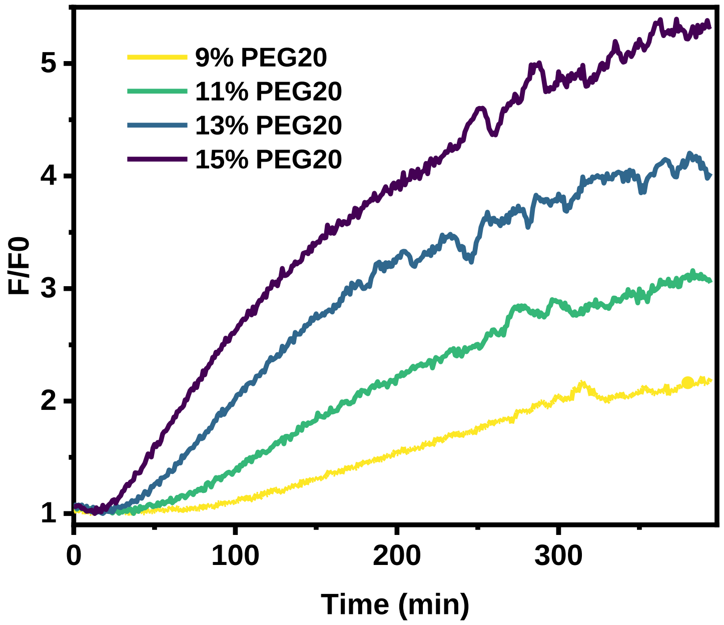


**Fig S6**. Thioflavin-T fluorescence intensity measurements with time after incubating 150 μM SUMO1 with different concentrations of PEG. The excitation wavelength used is 440 nm.

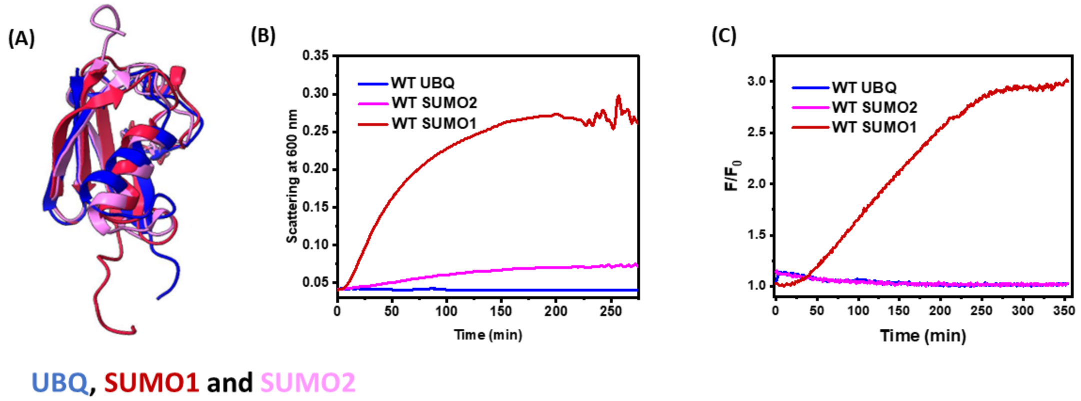


**Fig S7.** (A) Structural overlap of Ubiquitin (blue), SUMO1 (red) and SUMO2 (pink). (B) Comparison of change in scattering at 600 nm with time, after mixing 120 µM protein with 15% PEG. (C) Comparison of change in ThT fluorescence at 485 nm with time, after mixing 120 µM protein with 15% PEG, excitation wavelength used is 440 nm.

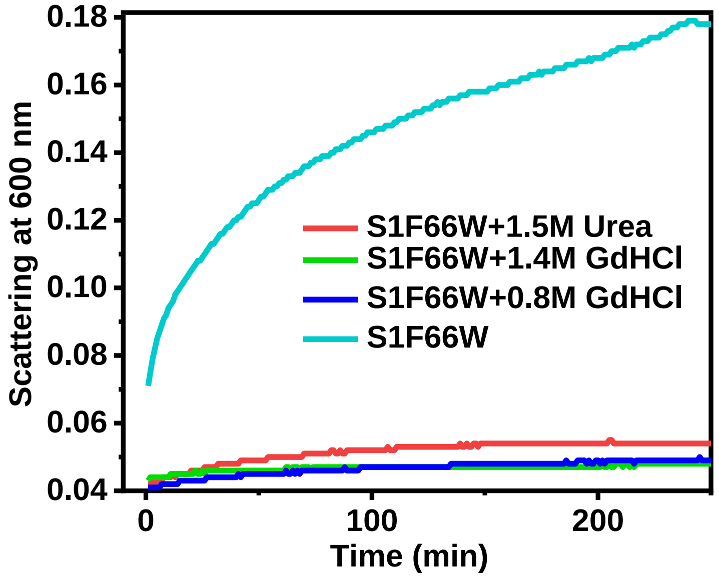


**Fig S8.** Comparison of scattering at 600 nm with time after mixing 120 μM of S1F66W containing no denaturant (cyan), 0.8 M (blue) and 1.4 M (green) GdHCl and 1.5 M Urea (red) with 15% PEG.

**Table S1**. Table for the lifetime components obtained by fitting the time-resolved experimental data in Figure 3B.

| sample | $\tau_{avg}$(ns) | $\alpha$1 | $\tau$1(ns) | $\alpha$2 | $\tau$2 (ns) | $\alpha$3 | $\tau$3 (ns) | $\chi^2$ |
|---|---|---|---|---|---|---|---|---|
| **S1F66W in PEG** | **0.54** | **0.66** | **0.13** | **0.31** | **0.32** | **0.03** | **1.29** | **0.95** |
| **S1F66W aggregates** | **1.57** | **0.66** | **0.14** | **0.19** | **0.52** | **0.15** | **2.26** | **0.95** |
| **S1F66W supernatant** | **0.57** | **0.61** | **0.15** | **0.35** | **0.33** | **0.04** | **1.61** | **0.96** |
